# Executable Simulation Model of the Liver

**DOI:** 10.1101/2020.01.04.894873

**Authors:** Matthias König

**Author notes:** **Correspondence**: Matthias König.

## Abstract

To address the issue of reproducibility in computational modeling we developed the concept of an executable simulation model (EXSIMO). An EXSIMO combines model, data and code with the execution environment to run the computational analysis in an automated manner using tools from software engineering. Key components are i) models, data and code for the computational analysis; ii) tests for models, data and code; and iii) an automation layer to run tests and execute the analysis. An EXSIMO combines version control, model, data, units, annotations, analysis, reports, execution environment, testing, continuous integration and release. We applied the concept to perform a replication study of a computational analysis of hepatic glucose metabolism in the liver. The corresponding EXSIMO is available from https://github.com/matthiaskoenig/exsimo.

## Introduction

We face a crisis in reproducibility [1], where it is impossible to believe most of the computational results shown in conferences and papers [2]. Recent replication efforts [3, 4] and theoretical considerations indicate that most published research findings in biomedical research are wrong [5].

A cornerstone of science is the possibility to critically assess the correctness of scientific claims and conclusions drawn by other scientists [6].

> *“An article about (computational) science in a scientific publication is not the scholarship itself, it is merely advertising of the scholarship. The actual scholarship is the complete … set of instructions and data which generated the figures.”*
>
> — — David Donoho

To be able to assess computational science we must be able to access the actual scholarship. But in the field of computational modeling in biology, most of the published quantitative models are lost because they are either not made available or they are insufficiently characterized [7]. Furthermore, for most studies neither the code nor data are accessible. Consequently, it is not possible to critically evaluate the correctness of the claims of most computational modeling analyses. This assessment has two main variants, reproducibility and replicability. “Reproducibility” means “running the same software on the same input data and obtaining the same results” [8] whereas “replicability” means “writing and then running new software based on the description of a computational model or method provided in the original publication, and obtaining results that are similar enough …” [8]. Reproducibility is a minimal standard, that something is reproducible does not imply that it is correct, the code most likely contains many bugs and methods may be poorly behaved. Replicability is much more stringent: Can someone repeat the experiment and get the same results [9].

An often overlooked fact increasing the challenge of reproducibility in computational modeling is that there will always be a next version of a computational analysis. The reasons are manifold, among them bugs in the model or analysis (or in libraries and software the analysis depends on), additional datasets to include in the analysis, additional simulation experiments to run with the model, model modifications to include additional processes, or changes in model parameters, to name a few.

To address the issue of reproducibility in versioned computational modeling projects we developed the concept of an executable simulation model (EXSIMO). We applied the concept to perform a replication study of a computational analysis of hepatic glucose metabolism in the liver [10].

## Results

To address the issue of reproducibility in the context of versioned computational modeling projects we developed the concept of an executable simulation model (EXSIMO). An EXSIMO combines model, data and code with the execution environment to run the computational analysis in an automated manner using tools from software engineering (Figure 1 and Figure 2). Key components are i) models, data and code for the computational analysis; ii) tests for models, data and code; and iii) an automation layer to run tests and execute the analysis. To demonstrate the concept an example EXSIMO based on a model of glucose metabolism in the liver (Figure 3) [10] was created and used to perform a replication study of the original work (Figure 4). In the following we walk through the different aspects of an EXSIMO using the example.

**Figure 1.**
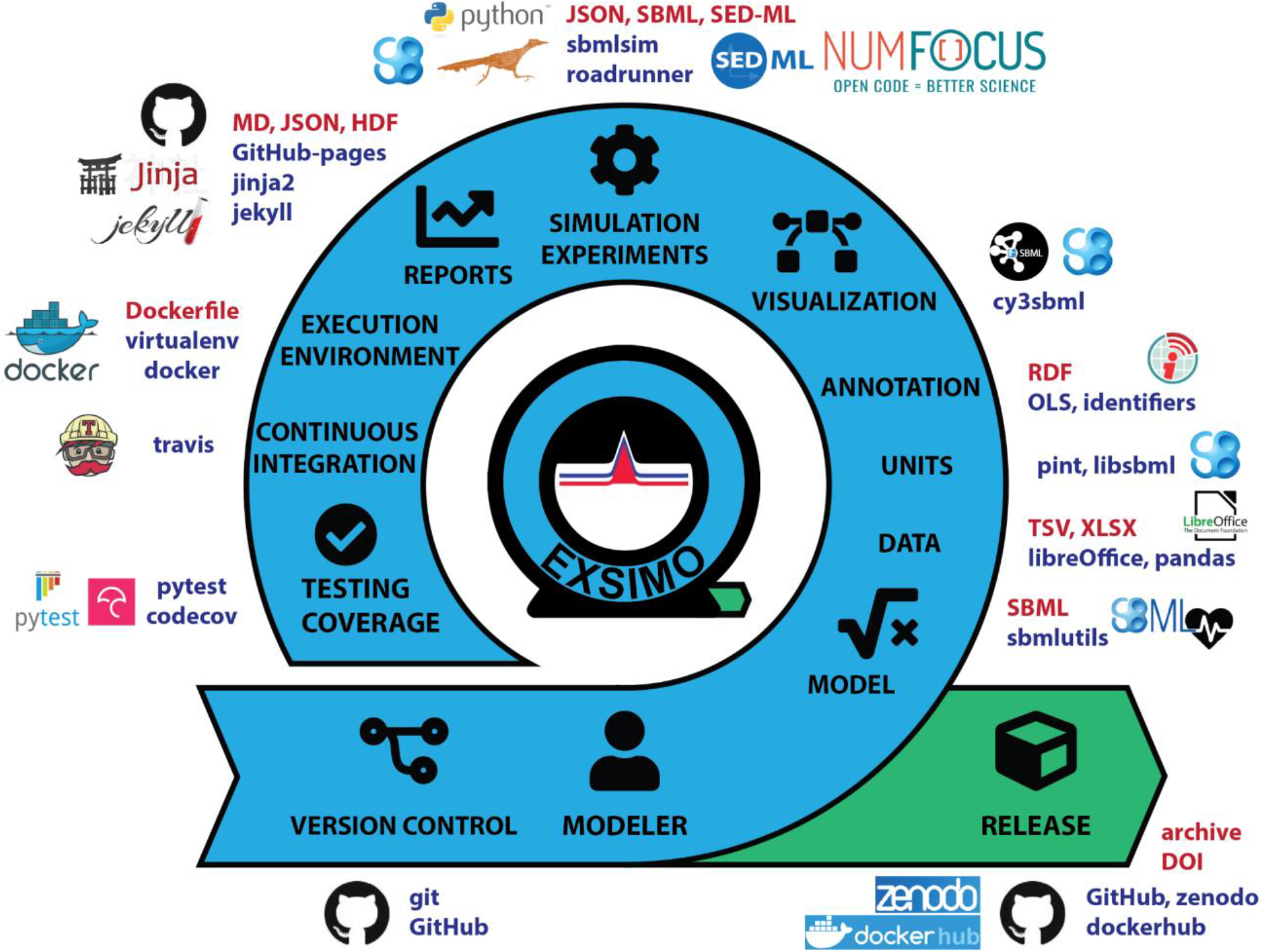
EXecutable SImulation MOdel (EXSIMO). An executable simulation model (EXSIMO) applies the tools and methods from software engineering to create high-quality reproducible model versions. Essential parts are tests to check model quality, datasets to evaluate model performance, and the definition of simulation experiments encoding the analysis performed with the model. All steps for testing and execution of the simulation model are automated within continuous integration. Formats used in the example EXSIMO are depicted in red, tools in blue. All formats are open standards, all tools are open-source and freely available for academic use.

### Version Control

> *“Talk is cheap. Show me the code.”*
>
> — — Linus Torvalds

Computational models and the corresponding analysis are continuously changing. Availability of all resources in the correct versions, i.e. models, data and code, is a prerequisite to reproduce the analysis. To keep track of these changes, EXSIMOs are built on top of version control. In our example we take advantage of the features of git and GitHub (https://github.com/matthiaskoenig/exsimo, Figure 2H) [11]. By using version control we enable collaborative work (merging changes), managing different versions (creating branches), tracking of changes (analyzing diffs), reuse (forking the repository), and versioning (using tags) of an EXSIMO. By using GitHub important features of managing a software project become available for an EXSIMO, e.g., code reviews, issue tracking, releases, team organization, and an automation layer via commit hocks, GitHub integrations and GitHub actions. Especially the release feature and automation are used heavily in an EXSIMO to automatically trigger testing, reporting and creation of release artifacts. Hosting code on publicly available repositories solves the code availability issue in most cases. Mangul et al. showed that software tool URLs directing the reader to online repositories have a high rate of accessibility; 99% of the links to GitHub and 96% of the links to SourceForge are accessible, while only 72% of links hosted elsewhere are accessible [12].

**Figure 2-.**
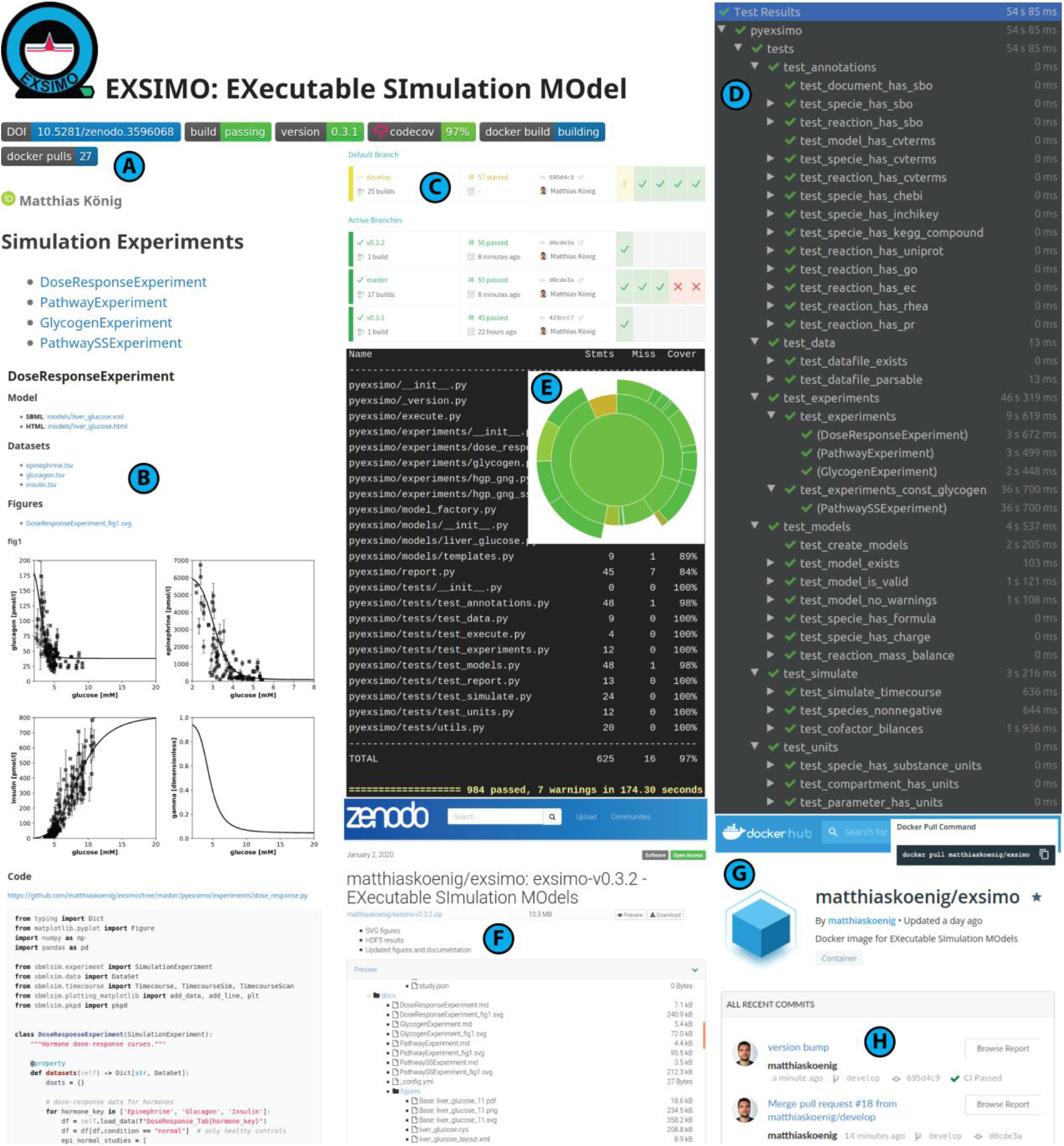
EXSIMO outputs. A) Automatically generated report for master branch (GitHub-pages serve markdown at https://matthiaskoenig.github.io/exsimo/). Report contains information on zenodo DOI, build status, release version, code coverage, corresponding docker image and executed simulation experiments. B) The reports for individual simulation experiments contain all SBML models, datasets, figures and code. The DoseResponseExperiment creates Figure 4A-D, PathwayExperiment Figure 4E-H, GlycogenExperiment Figure 4I-J, PathwaySSExperiment Figure 3K-M. C) Continuous integration with travis (https://travis-ci.org/matthiaskoenig/exsimo). After every commit all tests are executed including running the complete analysis. D) Overview over test functionality (984 tests are run in v0.3.2). E) Coverage of code by tests (https://codecov.io/gh/matthiaskoenig/exsimo). F) Zenodo release. Code with all results is packaged in a downloadable archive with DOI (https://doi.org/10.5281/zenodo.3596068). G) Tested execution environments are available as docker image from dockerhub (https://hub.docker.com/r/matthiaskoenig/exsimo). H) Code and issues are managed on GitHub with actions triggered by commits on respective branches (https://github.com/matthiaskoenig/exsimo). All results correspond to [26].

### Model

> *“A smart model is a good model.” —*
>
> — Tyra Banks

The computational model is a central piece of an EXSIMO. To ensure reproducibility and exchangeability of the models they must be encoded in a machine-readable standard exchange format. The de facto standard for model representation is SBML (Systems Biology Markup Language) [13, 14] and used as model format in the example. The model of hepatic glucose metabolism (Figure 3) is a metabolic pathway model based on ordinary differential equations (ODE) encoded in SBML using sbmlutils [15]. The model generation code is part of the repository and the SBML is generated on the fly during the analysis and in testing. Units and annotations (see below) are key meta-data smartening the model. All models are validated with libsbml [16].

**Figure 3-.**
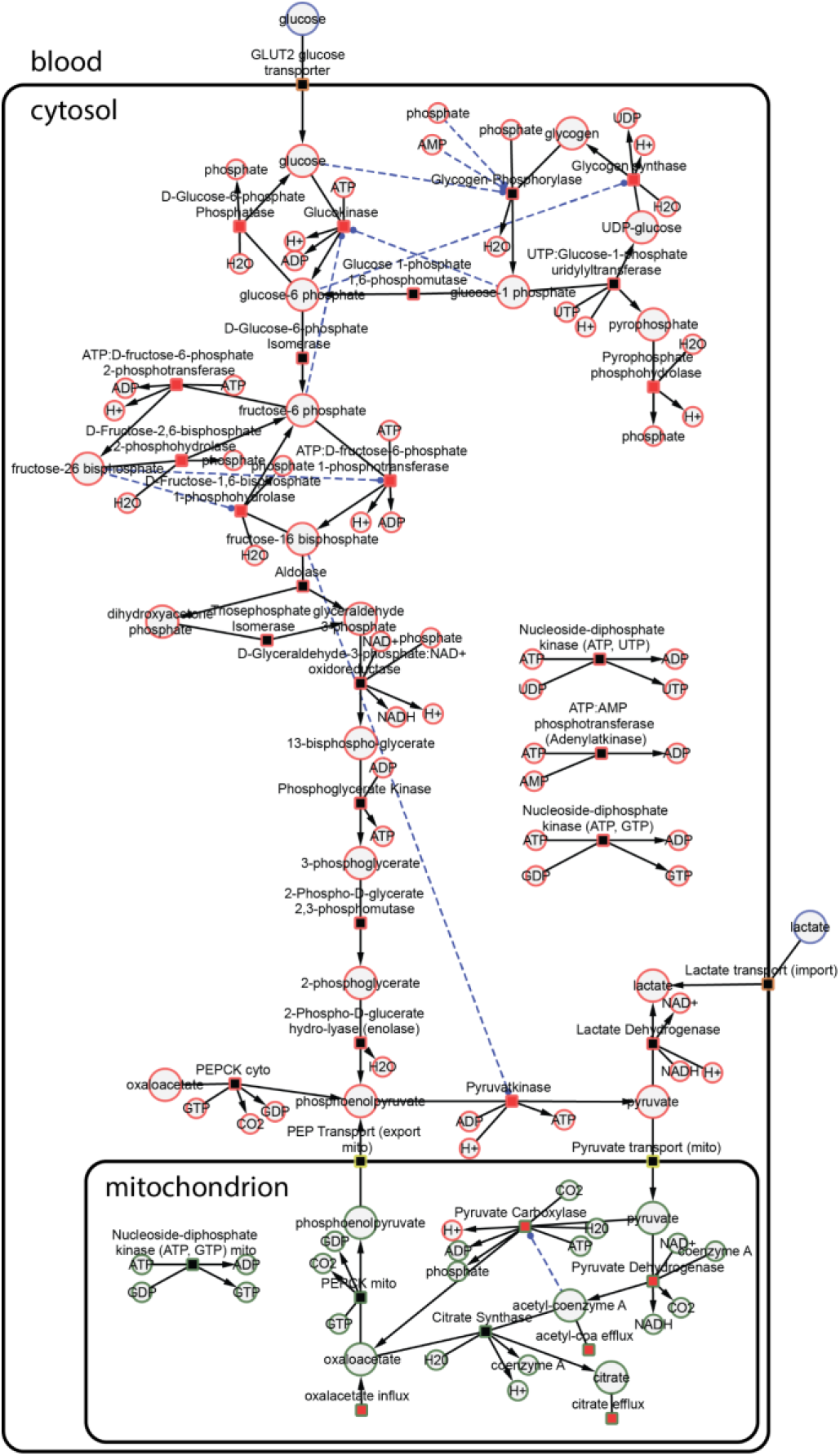
Model of Human hepatic glucose metabolism. The model was encoded in SBML using sbmlutils [15]. Model generation code and annotated SBML are available from https://github.com/matthiaskoenig/exsimo. Visualization was generated from SBML using cy3sbml [32, 33]. The biological description of the model and its components is available in [10].

### Data

> *“It is a capital mistake to theorize before one has data.”*
>
> — — Sherlock Holmes

Datasets are a crucial asset for parametrizing a model and for evaluating model predictions.

Data enables the reuse and modification of models, because the updated model can be fitted and its performance can be evaluated. A model without corresponding data sets is not very useful. In the example EXSIMO all experimental data were digitized from published figures and tables. Datasets are provided as Excel files with corresponding TSVs directly accessible from the reports. All data sources are described in [10] and the identical datasets were used in this replication study.

### Units

> *“Well, I use the metric system, It’s the only way to get really exact numbers.”*
>
> — — Catherynne M. Valente

Units are much more than meta-data, because equations only become meaningful if their units are clear [17]. A mathematical model without units is not a model. Therefore, an important part of an EXSIMO is to clearly define units for all model components. Due to complete unit information all model equations can automatically be unit validated (using libsbml model validation [16]). In addition units are annotated for all datasets used in the analysis. Only if units are specified on the model and datasets it is possible to ensure correct comparison of model predictions with data (especially because model units could change). A major advantage of units are automatic unit conversions in our example handled by pint within sbmlsim [18].

### Annotations

> *“The description is not the described … The thought is not the thing.”*
>
> — — Jiddu Krishnamurti

Semantic annotations are meta-data making models and data useful. Semantic annotations describe the computational or biological meaning of models and data via machine-readable links to knowledge resource terms. These annotations help to find models and datasets, accelerate model composition and enable knowledge integration between models and experimental data [19]. Within the EXSIMO it is tested that the minimum information requested in the annotation of biochemical models (MIRIAM [7]) is fulfilled. The example model is annotated using resolvable identifiers and resources from identifiers.org [20]. Most model components are annotated with sbo terms, species with inchikey, chebi, and kegg.compound terms, reactions with rhea, uniprot, go and ec-code terms. In addition, charge and chemical formula are stored for all species which allows to perform tests of mass and charge balance on all reactions.

### Simulation Experiments

> *“A computer lets you make more mistakes faster than any invention in human history, with the possible exceptions of handguns and tequila.”*
>
> — — Mitch Radcliffe

In an EXSIMO the actual computational analysis is encoded in the form of simulation experiments (similar to SED-ML [21, 22]). Every one of this experiments defines the necessary datasets, simulation tasks, and data processing to create output figures for a given question. Specifically, we performed a replication study of König et al., 2012 [10] with results depicted in (Figure 3 and Figure 4). The individual parts of the replication were encoded in four simulation experiments: i) DoseResponseExperiment (Figure 4A-D), PathwayExperiment (Figure 4E-H), GlycogenExperiment (Figure 4I-J) and PathwaySSExperiment (Figure 4K-M).

**Figure 4-.**
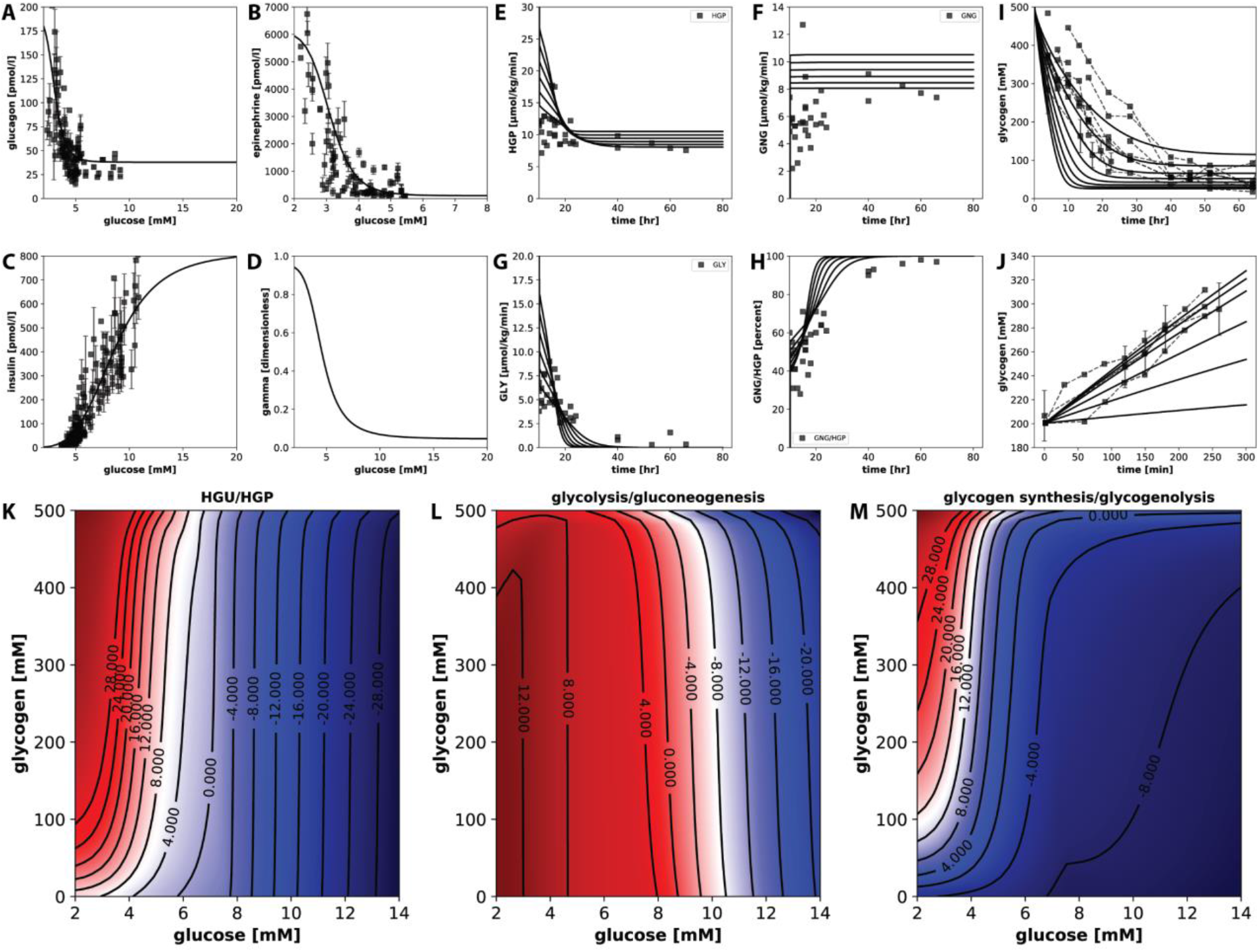
Results replication study. Results from [10] are replicated via simulation experiments. The report including SBML models, datasets, figures and code is available from https://matthiaskoenig.github.io/exsimo/). Panels A-D correspond to Figure 2A-D (hormonal dose-response curve) in [10], panels E-H to Figure 3A-D (time course of hepatic glucose production (HGP), gluconeogenesis (GNG), glycogenolysis (GLY) and contribution of gluconeogenesis to hepatic glucose production (GNG/HGP) under varying glucose concentrations), panels I-J to Figure 4A-B (time course glycogenolysis and glycogen synthesis under varying glucose concentrations), panels K,L,M to Figure 5A,C,D (steady state scan at various glucose and glycogen concentrations of hepatic glucose utilization (HGU), hepatic glucose production (HGP), glycolysis, gluconeogenesis, glycogen synthesis and glycogenolysis). For biological interpretation see [10]. All results correspond to [26]. Data are means ± SD.

The original analysis was implemented in Matlab with model equations directly in the code simulated with ode15s (no working SBML was available). In this replication an annotated and unit-validated SBML was created. The individual analyses were reverse-engineered from the original figures and encoded in python as simulation experiments using sbmlsim [18]. When things were insufficiently described in the publication we used the original source code to clarify issues (https://github.com/matthiaskoenig/glucose-model). Replication simulations were performed using roadrunner [23].

An important outcome based on this replication is that the original model was incorrectly implemented in Matlab. Only by translating the ODES to SBML the issues became apparent. Specifically, the glycogen pathway was scaled in the ODEs in a manner which was not compatible with a species-reaction description. Without any checks and tests on the ODE system, nor tests on model structure in Matlab these errors went undetected. As a consequence of this bugs numerical differences between the original paper [10] and this replication exist (specifically in the glycogen reactions). Replication is defined as repeating the same experiments with different methods and getting similar results. All biological relevant model behavior could be replicated with very similar outcomes to the original work, i.e. i) dose-response curve of hormones; ii) time-dependent hepatic glucose production (HGP) via gluconeogenesis (GNG) and glycolysis (GLY) as well as the ratio GNG/HGP; iii) time-dependent glycogenolysis and glycogen synthesis under various blood glucose concentrations; and iv) steady state scan of HGP, GLY and GNG under varying blood glucose and glycogen concentrations. Especially important the correct switching points of the pathways are replicated.

By encoding the model in SBML and adding thorough model tests on top of the SBML validation we could ensure correct model behavior in EXSIMO.

### Reports

> *“Numbers have an important story to tell. They rely on you to give them a clear and convincing voice.”*
>
> — — Stephen Few

An important aspect of an EXSIMO is a human readable report and summary of the computational analysis (Figure 2A-B) available from https://matthiaskoenig.github.io/exsimo/. This includes all assets and information for the respective simulation experiments, i.e. models, datasets, figures and executed code. These reports are generated automatically from commits to the master branch (GitHub-pages serve markdown created in the analysis). The report includes information on zenodo DOI, build status, release version, code coverage, corresponding docker image and executed simulation experiments. Simulation results are stored in compressed HDF5 for all executed simulations (but not tracked in git due to their large file sizes).

### Execution Environment

> *“I don’t care if it works on your machine! We are not shipping your machine!”*
>
> — — Vidiu Platon

To ensure reproducibility of a computational analysis the execution environment must be provided as an artifact. Within the EXSIMO, all code is provided as an easily installable package (python package installable with pip with dependencies recorded using a requirements file). This allows to setup a python virtual environment for the computational analysis via a single line of code for instance in testing. Unfortunately, this information is not sufficient to guarantee reproducibility because unpinned package versions exist in the package dependency tree (not all versions are specified exactly) and package builds depend on system libraries. For most practical aspects such an environment can be considered reproducible, but many corner cases exist which can break the computational analysis. To solve this issue we distribute in addition the execution environment as docker images (https://hub.docker.com/r/matthiaskoenig/exsimo, Figure 2G) with builds of the images triggered by GitHub commits. As part of the image creation the complete tests and analysis are executed, thereby ensuring the analysis can be run in the container.

### Testing

> *“If debugging is the process of removing bugs, then programming must be the process of putting them in.”*
>
> — — Edsger Dijkstra

Unit tests are used to ensure model quality and that all simulation experiments can be executed correctly. An overview over tested functionality is depicted in Figure 2D. Tests check for example successful SBML generation, validity of the model, existence and correctness of annotations and units, mass and charge balance on reactions or cofactor balances. The 984 tests cover 97% of the code (Figure 2E, https://codecov.io/gh/matthiaskoenig/exsimo). Similar to the unit and integration testing practices in software engineering, example simulations with a description of the expected results allows to verify that the EXSIMO was successfully installed and works correctly [12]. A similar strategy of using unit tests to ensure model quality has been applied in other modeling projects, e.g., for genome-scale metabolic models in memote [24] or in the OpenWorm project [25].

### Continuous Integration

> *“The most powerful tool we have as developers is automation.”*
>
> — — Scott Hanselman

Continuous integration (CI) is the practice of merging code to a shared main code line, i.e. local changes to the develop branch or the develop branch to the master branch. CI is used in an EXSIMO in combination with automated unit testing, i.e. after every commit all tests are executed including running the complete analysis (Figure 2D). Within the example we use travis (Figure 2C, https://travis-ci.org/matthiaskoenig/exsimo) integrated with GitHub running the tests in the defined execution environments.

### Release

> *“It is easier to do a job right than to explain why you didn’t.”*
>
> — — Martin Van Buren

The final step is to create a release for a given version. This is handled via GitHub releases from the master branch, which automatically trigger creation of the reports, upload to zenodo and building of the docker image. Zenodo provides a downloadable archive containing model, data, code and results which can be referenced uniquely via a DOI (Figure 2F, [26]). Importantly, as part of every release the computational analysis is reproduced and tested on three different systems: i) in a local virtual environment; ii) in an ubuntu 18.04 virtual environment during continuous integration by travis; iii) in a python 3.6 docker image during building the docker image by docker hub (Figure 2G).

## Discussion

Within this work we presented the concept of an executable simulation model (EXSIMO) with reproducibility by design and demonstrated it by performing a replication study of a model of glucose metabolism in the liver [10].

EXSIMOs enable continuous and rapid development of models and computational modeling analysis via a workflow closely mimicking how models are developed in the praxis. The presented example EXSIMO is one possible implementation of an executable simulation model. For the individual parts alternative tools can be chosen. For instance, one could use mercurial with sourceforge for version control, use CellML as model description language, encode data in JSON, write analysis code in R, create reports in latex, store the execution environment as virtual machine, use Jenkins continuous integration, and release to Figshare. The concept remains, only the tooling changes.

Reproducibility in computational modeling builds on top of community-based information standards (COMBINE) [27] and FAIR data [28], both integral parts of an EXSIMO. Computational models are represented in SBML [13, 14], simulation experiments are compatible to SED-ML [22], released archives closely mimic COMBINE archives [29] and annotations utilize BioModels.net Qualifiers, identifiers.org URIs [20] and SBO (Systems Biology Ontology) terms. Both, minimal information standards for representation of models (MIRIAM) [7] and simulation experiments (MIASE) [30] are fulfilled. All data is findable, accessible, interoperable and reusable (FAIR). Future work will further improve support for these community standards in the context of executable simulation models.

At the moment it is impossible to believe most of the computational results shown in conferences and papers [2]. “An article about (computational) science in a scientific publication is not the scholarship itself, it is merely advertising of the scholarship. The actual scholarship is the complete set of instructions and data which generated the figures.” [31]. Most papers with computational models do neither provide the models in a machine-readable format, the data used to fit or evaluate the model, nor the code underlying the analysis. One may ask, how were reviewers able to evaluate the actual scholarship in all these publications? Without the ability to critically assess the correctness of scientific claims, the scientific methods fails. So next time you review an article or grant proposal stop for a second and ask: Is this advertisement or actual scholarship?We urgently need a change in culture how computational modeling studies are published, reviewed and evaluated by the community. A rigorous standardized approach is needed to examine software tools prior to publication [12]. With computational modeling studies being small software projects, similar rigorous approaches must be applied. For a publication on computational modeling we should not accept less than complete linked and executable code and data. Disseminating a computational modeling analysis as an executable simulation model with reproducibility by design could be one approach to restore credibility and trust in computational modeling.

## Author Contributions

MK designed the study, performed the analysis, and wrote the manuscript.

## Acknowledgements

MK is supported by the Federal Ministry of Education and Research (BMBF, Germany) within the research network Systems Medicine of the Liver (LiSyM, grant number 031L0054).

